# The evolutionary dynamics of Oropouche Virus (OROV) in South America

**DOI:** 10.1101/682559

**Authors:** Bernardo Gutierrez, Emma Wise, Steven Pullan, Christopher Logue, Thomas A. Bowden, Gabriel Trueba, Marcio Nunes, Nuno R. Faria, Oliver G. Pybus

## Abstract

The Amazon basin is host to numerous arthropod-borne viral pathogens that cause febrile disease in humans. Among these, *Oropouche orthobunyavirus* (OROV) is a relatively understudied member of the Peribunyavirales that causes periodic outbreaks in human populations in Brazil and other South American countries. Although several studies have described the genetic diversity of the virus, the evolutionary processes that shape the viral genome remain poorly understood. Here we present a comprehensive study of the genomic dynamics of OROV that encompasses phylogenetic analysis, evolutionary rate estimates, inference of natural selective pressures, recombination and reassortment, and structural analysis of OROV variants. Our study includes all available published sequences, as well as a set of new OROV genomes sequences obtained from patients in Ecuador, representing the first set of viral genomes from this country. Our results show that differing evolutionary processes on the three segments that encompass the viral genome lead to variable evolutionary rates and TMRCAs that could be explained by cryptic reassortment. We also present the discovery of previously unobserved putative N-linked glycosylation sites, and codons which evolve under positive selection on the viral surface proteins, and discuss the potential role of these features in the evolution of the virus through a combined phylogenetic and structural approach.

## Introduction

The Bunyavirales is a highly diverse order of viruses that include multiple emerging human pathogens. Within this order, the *Orthobunyavirus* genus (family: *Peribunyaviridae*) includes many arthropod-borne virus species that have been associated with disease in humans (1). The severity of disease varies and ranges from self-limiting febrile illness to encephalitis and haemorrhagic fever (2). One prominent member of this genus is *Oropouche orthobunyavirus* (OROV), a common causal agent of febrile disease in the Amazon basin (3) and a potential candidate for future emerging epidemics in the region and elsewhere (4).

Since its first description in 1961 in Trinidad and Tobago (5), OROV has caused several outbreaks and sporadic infections in the Brazilian Amazon, particularly in the states of Pará, Amapá, Rondônia, Maranhão, Acre, Amazonas, Minas Gerais, Mato Grosso, and Tocantins (6–9), and evidence suggests the circulation of OROV in other central Brazilian states (10). Since the late 1980s, additional cases and outbreaks of OROV have been reported in Panama, Peru (11–13) and, more recently, Ecuador (14). The virus has been isolated from sloths (*Bradypus trydactilus*) (3) and marmosets (*Callithrix sp.*) (15) as well as from invertebrates such as *Culex p. quinquefasciatus* (9) and *Ochlerotatus serratus* mosquitoes and *Culicoides paraensis* biting midges (3). It has been proposed that OROV has two distinct transmission cycles (16). In the sylvatic transmission cycle, wild mammals such as sloths (*B. trydactilus*) and primates (*Allouatta caraya*, *Callithrix penicillata*) serve as hosts for the virus, which is transmitted by vector species that are common in rural areas (*Aedes serratus* and *Coquillettidia venezuelensis* have been proposed as vectors in the sylvatic cycle). This sylvatic cycle may possibly include other vertebrate hosts, including wild birds and rodents (such as *Proechimys sp.*). In contrast, the urban transmission cycle involves human-to-human transmission mediated by the biting midge *C. paraensis* (16) and can be facilitated by anthropogenic disturbance of forest areas. OROV produces an acute febrile illness in humans, characterised by unspecific symptoms such as fever, chills, headaches, myalgia and joint pain (17) and is therefore difficult to diagnose and differentiate from other co-occurring infectious diseases such as leptospirosis, dengue fever, Venezuelan Equine Encephalitis (VEE), malaria, and rickettsial and *Coxiella* infections (18)

The genome of OROV is typical of the *Orthobunyaviruses* and is comprised of three negative-sense RNA segments of different sizes: a large segment (L) that encodes a RNA dependent RNA polymerase, a medium (M) segment that encodes a membrane polyprotein comprised of three main components (the Gn, NSm and Gc proteins), and a small (S) segment that encodes the structural nucleocapsid protein N and a second non-structural protein, Ns, in an overlapping reading frame (2). The Gn and Gc proteins determine the architecture of the viral particle; CryoEM studies show that the membrane proteins of Bunyamwera virus (BUNV) form a tripodal structure (19) and the crystal structures of the N-terminus of Gc suggest that this trimeric assembly may be conserved across various Orthobunyaviruses (20). The segmented nature of the OROV genome enhances the likelihood of genome reassortment events, a phenomenon that has been explored more generally in the Bunyaviruses (2) and has been associated, in some cases, with a dramatic increase in disease severity, such as with Ngari virus, a reassortant between Bunyamwera virus (BUNV) and Batai virus (BATV) (21). Furthermore, reassortment appears to play an important role in the evolution and emergence of new viruses within OROV and related species, as has been described for the reassortant species Jatobal virus (JATV) in Brazil (22), Iquitos virus (IQTV) in Peru (23) and Madre de Dios virus (MDDV) in Venezuela (24).

Despite a heightened interest in OROV, reflected by an increasing number of publications covering the topic in recent years (16), many aspects of OROV history, biology and ecology remain poorly understood. Although the evolution of the virus has been previously studied using standard phylogenetic methods (3, 7, 8, 25, 26), a comprehensive analysis of the molecular evolution of OROV that incorporates the complete genetic diversity of the virus and compares the evolutionary histories of the genome segments has not yet been undertaken. Here we present the results of an in-depth analysis of the OROV genome that includes the genomes of 6 new OROV isolates sampled in Ecuador and explores the differences between the evolutionary histories of the three viral segments through comparative phylogenetics. We evaluate the temporal signal of each segment, estimate their dates of origin and rates of evolution, and test for recombination and reassortment. Furthermore, we combine phylogenetic tests for selection on the viral genome with structural mapping and N-linked glycosylation sequon analysis to determine the factors that can drive the evolution of the viral genome and explain the observed differences between segments.

## Methods

### Sampling and sequencing of OROV isolates from Ecuador

Six strains of OROV were isolated independently via one passage in Vero cells, from febrile patient sera sampled in 2016 in the Esmeraldas region of Ecuador. RNA was extracted and six complete genome sequences were generated using a metagenomic approach as described previously (14). Briefly, cDNA was prepared using a Sequence Independent Single Primer Amplification (SISPA) approach (27), cDNA sequencing libraries were prepared using the Nextera XT Kit (Illumina), and sequencing was performed on a MiSeq (Illumina) using 150 nucleotide paired-end runs. Reads were mapped to reference sequences using BWA MEM (28). The final consensus sequences were generated using Quasibam (29) The study was approved by the bioethics committee of Universidad San Francisco de Quito, and all patients provided written consent indicating that they agreed for their samples to be tested for additional pathogens.

### Data sets

In addition to the Ecuadorian sequences, we collected all available sequences for the three OROV genome segments from GenBank at NCBI. We then excluded all sequences that were shorter than a threshold segment length (specifically, <6500 nucleotides long for L, <4200 nucleotides long for M and <693 nucleotides long for S). These GenBank sequences represent the genetic diversity of OROV from outbreaks in Trinidad & Tobago, Panama, Peru and Brazil, and cover the complete sampled history of the virus from the mid 1950s to the late 2000s, with the latest outbreak in Brazil reported in 2009 (7). The data sets were compiled into individual alignments for each of the three viral genome segments (denoted L for long, M for medium and S for short).

Additional data for the OROV sequences included the sampling date (with varying precision of year, month or exact day, depending on the sequence), the identity of the host species (most samples were obtained from human patients, while a few were obtained from potential arthropod vectors and mammalian reservoir species), and the location of each sample (province of origin was used for the samples from Brazil and Peru, while only the country of origin was available for the Panama sequences; samples from Ecuador and Trinidad & Tobago were obtained from a single location). No exact sampling date is available for the Ecuadorian sequences, but sampling was performed between the months of April and May of 2016 (Sully Márquez, Pers. Comm.).

Before sequence alignment, the untranscribed terminal repeats (UTRs) of each segment were removed and the open reading frame of each segment was identified (only the first ORF of the S segment was analysed). Sequences were then aligned using the MUSCLE algorithm, as implemented in Geneious 9.0.5 (30), and the alignments were manually checked and edited to ensure they were codon-aligned. The L alignment included 59 published sequences plus the six new sequences from Ecuador. The M segment alignment included 58 published sequences and the six Ecuadorian sequences. Finally, the S segment alignment included 143 published sequences plus the six Ecuadorian sequences.

### Molecular phylogenetics and molecular clock analyses

All three alignments were scanned for recombinant sequences using the suite of methods implemented in RDP4 (31). Sequences identified as recombinant (i.e. evidence for recombination was found by at least four methods) were removed from all further analyses (only two M segment sequences were excluded). For each alignment, a molecular phylogeny was estimated using the maximum likelihood (ML) approach implemented in RAxML 8.0 (32). ML trees were estimated using a General Time Reversible (GTR) substitution model with a Gamma-distribution model of among-site heterogeneity. Phylogenetic node support was assessed using a non-parametric bootstrap approach with 100 replicates. No outgroup sequences were used and all trees were midpoint rooted. A visual comparison of the tree topologies and bootstrap scores for each segment was used to evaluate possible OROV reassortment events during its evolutionary history.

To explore the rates of OROV molecular evolution and to evaluate the temporal signal in each OROV alignment, we correlated tip-to-root genetic distances in the ML tree against the sampling dates of the corresponding sequences, using the approach implemented in TempEst 1.5.1 (33). The regression plots were inspected visually to identify notable outliers and the linear correlation coefficient of the regression was used as a measure of the degree to which the sequences evolve in a clock-like manner.

We estimated the evolutionary history of OROV outbreaks in South America by constructing time-calibrated Maximum Clade Credibility (MCC) trees for each segment. Each MCC tree summarises a posterior distribution of phylogenies that were sampled using the Bayesian Markov Chain Monte Carlo (MCMC) approach implemented in BEAST 1.10.1 (34). Trees were sampled under a HKY substitution model, a Gamma-distribution model of among-site heterogeneity, a Bayesian Skyline tree prior (35), and a relaxed uncorrelated molecular clock (UCLD) (36) model.

Our ML and TempEst analyses (above) revealed two monophyletic lineages in the M segment phylogeny, one with weak temporal signal (correlation coefficient=0.66) and one with stronger signal (correlation coefficient=0.94) (see *Results* and Fig. S1). Therefore, the molecular clock analysis of the M segment was performed in a two-step process. First, we estimated a time-calibrated tree separately for the two group M lineages and noted the estimated time of the most recent common ancestor (TMRCA) of the lineage with stronger temporal signal (Lineage 2). We then performed a second analysis of the whole group M phylogeny, in which this TMCRA estimate was used as an additional calibration point (i.e. as an informative prior on the date of the common ancestor of Lineage 2). Similarly, mean evolutionary rates were co-estimated with phylogeny for each of the three segments using the UCLD clock model and subsequently compared. We further repeated this analysis separately for M segment lineages 1 and 2.

### Selection Analyses

The ORF alignments of the S and L segments, as well as those of the individual proteins of the M segment ORF (Gn, NSm and Gc) were tested for evidence of natural selection using the various methods implemented in HyPhy (37), implemented by the Datamonkey 2.0 server (38). Statistical tests were used to evaluate evidence for both pervasive and episodic selection in the OROV genome, and to evaluate selection across whole genes as well as at specific codons. Analyses of pervasive selection were performed using three methods: (i) the Branch-site Unrestricted Statistical Test for Episodic Diversification (BUSTED) method, which tests for gene-wise selection and estimates a mean *dN*/*dS* ratio for each gene (39), (ii) the Single-Likelihood Ancestor Counting (SLAC) method, and (iii) the Fixed Effects Likelihood (FEL) method; the latter two approaches were used to infer specific codons under adaptive selection (40). Episodic selection was evaluated using two methods: (i) the Mixed Effects Model of Evolution (MEME), which identifies individual codons under positive selection (41), and (ii) a branch-site model implemented in aBSREL (42), which was used to search for phylogenetic branches under selection. We identified sites as being positively selected with a critical value of p=0.05. Relative codon positions were numbered according to the OROV reference strain BeAn19991.

### Structural analysis of OROV proteins and sites under selection

The availability of high-resolution protein structures for the OROV proteins is limited. We therefore focused efforts on the OROV glycoprotein Gc because (i) N-terminal regions for a number of orthobunyaviral Gc glycoproteins (including the head domain of OROV Gc, PDB ID 6H3X), have been structurally elucidated (20) and (ii) it shows an increased number of sites under positive selection compared to the other OROV proteins (see Results). The second component of the OROV glycoprotein, Gn, was not analysed here because no adequate atomic resolution structure is currently available. For Gc, N-linked glycosylation sites were predicted for each virus sequence using the NetNGlyc 1.0 server (www.cbs.dtu.dk/services/NetNGlyc/), which identifies NXT/S amino acid motifs (where X is any amino acid except proline). The sequence evolution of these motifs was subsequently mapped onto the M segment phylogeny using a maximum parsimony approach. To enable structure-based mapping of sites under selection, we created a composite molecular model that encompasses the structurally elucidated N-terminal head of OROV Gc (PDB ID 6H3X) and the stem domains from the closely related Schmallenberg virus (SBV) (PDB ID 6H3S) (20) through sequence-independent structural alignments in PyMOL (43). As there are no known structures of the orthobunyaviral C-terminal core, this region was omitted from the structural analysis.

We explored the evolutionary history of molecular adaptation that has shaped the different domains of the Gc protein by estimating dN/dS ratios for each protein domain using the renaissance counting approach (44) implemented in BEAST 1.10.1. These estimates were independently performed for the full set of M sequences and for the two previously described lineages, using an empirical posterior sample of phylogenies comprising >100,000 trees. The 95% highest posterior distribution (HPD) credible intervals for the normalised dN and dS values were summarised using the *coda* package in R (the normalised values are the observed count of each substitution type, divided by the expected count of each substitution type under the substitution and evolutionary models and independent of the data).

## Results

### Molecular phylogenetics and molecular clock analyses

The estimated ML phylogenies for each OROV genome segment are shown in Figure 1 and reveal a pattern of spatial structure. In all three phylogenies (Fig 1a, b, c), our new sequences from Ecuador (yellow) cluster together in a single monophyletic group (with 100% bootstrap support) that is most closely related to OROV sequences from Peru (red), suggesting the Ecuadorian sequences represent an unreported outbreak in 2016 that is related to cases observed in Peru in the mid-to-late 2000s. The large (L) and medium (M) segment trees (Fig 1b, c) contain long internal branches and distinct lineages. Sequences from Panama (orange) are found in two distinct clusters, while Brazilian sequences (light blue) are split into two distinct groups. One Brazilian group corresponds to the 2009 OROV outbreak in the municipality of Mazagão (Amapá state) in northern Brazil (7), whilst the other contains sequences from isolates obtained between 1960 and 2006 from seven different states (Acre, Amapá, Amazonas, Maranhão, Minas Gerais, Pará and Rondônia) (45). Most of these clusters are supported by high bootstrap scores (>95%) but there is less consistent and sometimes limited support for phylogenetic structure *within* clusters. The OROV small (S) segment phylogeny (Fig 1a) contains more sequences but has less phylogenetic structure (i.e. internal branches are shorter and several basal nodes have weak bootstrap support). Sequences still tend to cluster according to sample location; there are 3 well supported clusters of sequences from Brazil or Brazil/Panama (Fig. 1). The Ecuadorian sequences are again found in a strongly supported cluster together with Peruvian isolate IQE7894.

**Figure 1.**
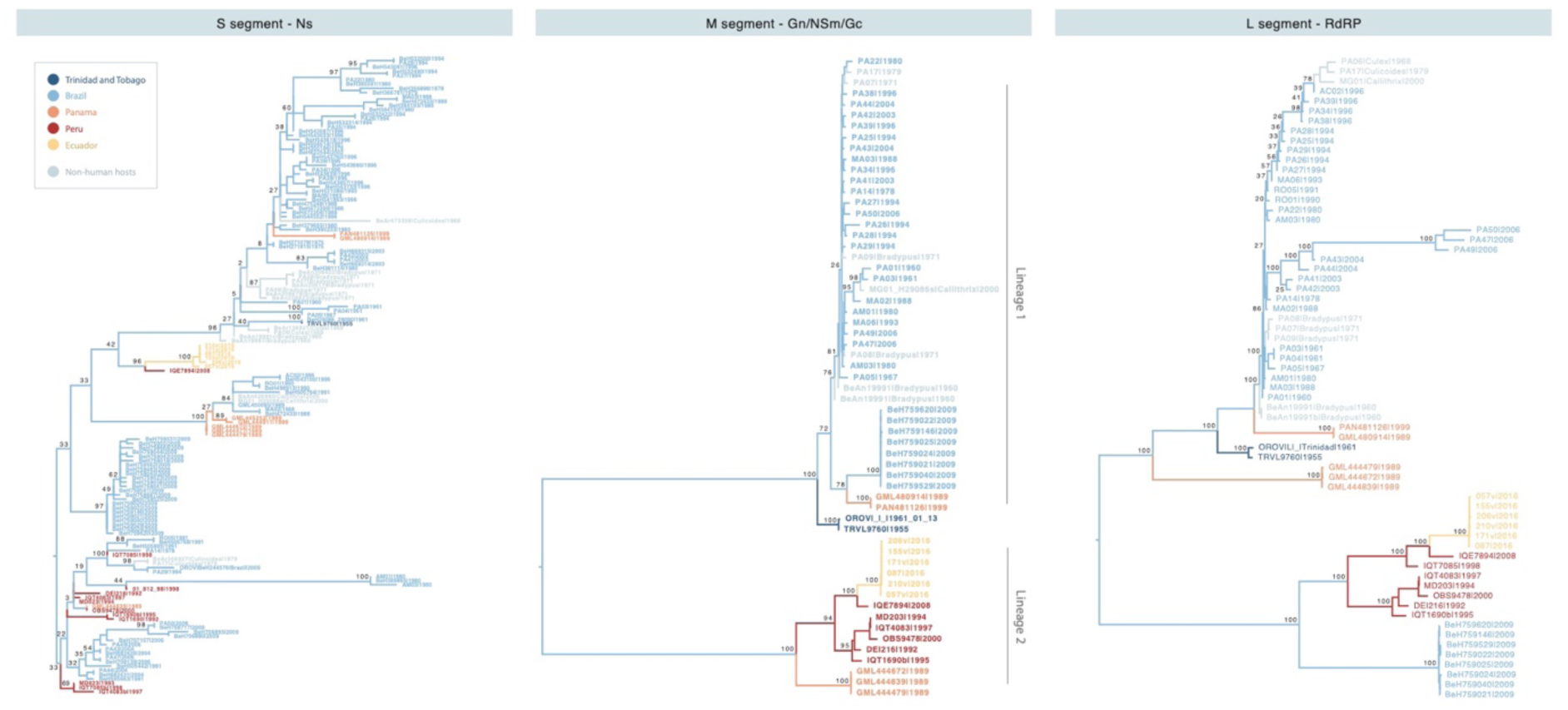
Maximum Likelihood trees for the small (S), medium (M) and large (L) genomic segments of the OROV genome. Tip labels and corresponding branches are colour coded by the country of origin for each sample, with grey samples indicating sequences obtained from non-human hosts and vectors (species indicated in the tip label). Node support from 100 bootstrap replicates is shown for the basal internal nodes of each tree.

The trees for the different segments show considerable topological differences, suggesting the possibility of reassortment events in the viral genome. Pairwise comparisons between segment phylogenies (Fig. S2) indicates that the L segment of the 2009 Mazagão outbreak sequences has arisen through a reassortment event. There are also widespread topological differences between the S segment and the L and M segments (Fig. S2), particularly for the Brazilian sequences (which cluster in robust monophyletic clades in the L and M segment trees but as multiple clusters in the S segment phylogeny). This suggests that OROV might reassort more frequently in the heart of the Amazon basin, possibly due to the co-circulation of viral lineages (many of which may be currently unsampled) and the co-infection of wild hosts in the Amazonian forest. However, the low node support for the S segment could also represent phylogenetic uncertainty rather than differing evolutionary history.

The evolutionary timescale for OROV in America, estimated using a Bayesian phylogenetic molecular clock approach, yielded highly variable dates for the most recent common ancestor (TMRCA) for each segment. The origins of the S and L segments can be traced back to the early to mid 20^th^ century, while the M segment shows long internal branches and a much older root, dating back to approximately 1675 (95% HPD: 1115.68-1938.77) (Fig. 2a). Despite the fact that the exact geographical location where OROV emerged remains unknown, the estimated TMRCA of the S segment is approximately 1940 (95% HPD: 1916.15-1955.00), while that of the L segment is slightly older, around 1921 (95% HPD: 1860.66-1955.00). It is also noteworthy that the M segment splits into two distinct lineages (Fig. 2a). Lineage 1 represents an eastern-central clade that includes the earliest OROV case in Trinidad and Tobago, sequences obtained from patients in Panama between the late 80s and late 90s, and the entirety of the Brazilian genetic diversity of the virus. Lineage 1 is estimated to have emerged around 1899 (95% HPD: 1801.20-1948.83). On the other hand, Lineage 2 contains sequences from western South America, including sequences from Panama (from the late 1980s), Peru and Ecuador, with an estimated origin around 1918 (95% HPD: 1806.05-1970.84). The estimated evolutionary rate of the M segment appears to be significantly lower than that of other segments (mean rate = 1.84×10^−3^ subst./site/year for segment S, 3.89×10^−4^ subst./site/year for segment M, and 1.61×10^−3^ subst./site/year for segment L). The evolutionary rates estimated separately for each lineage within the M segment phylogeny indicates that Lineage 2 evolves considerably faster than Lineage 1, which more closely resembles the overall M segment rate (mean rate =2.30×10^−4^ subst./site/year for Lineage 1, and 8.34×10^−3^ subst./site/year for Lineage 2) (Fig. 2b). However, due to the large credible regions for the Lineage 2 rate, we cannot conclude that it evolves faster than the other segments.

**Figure 2.**
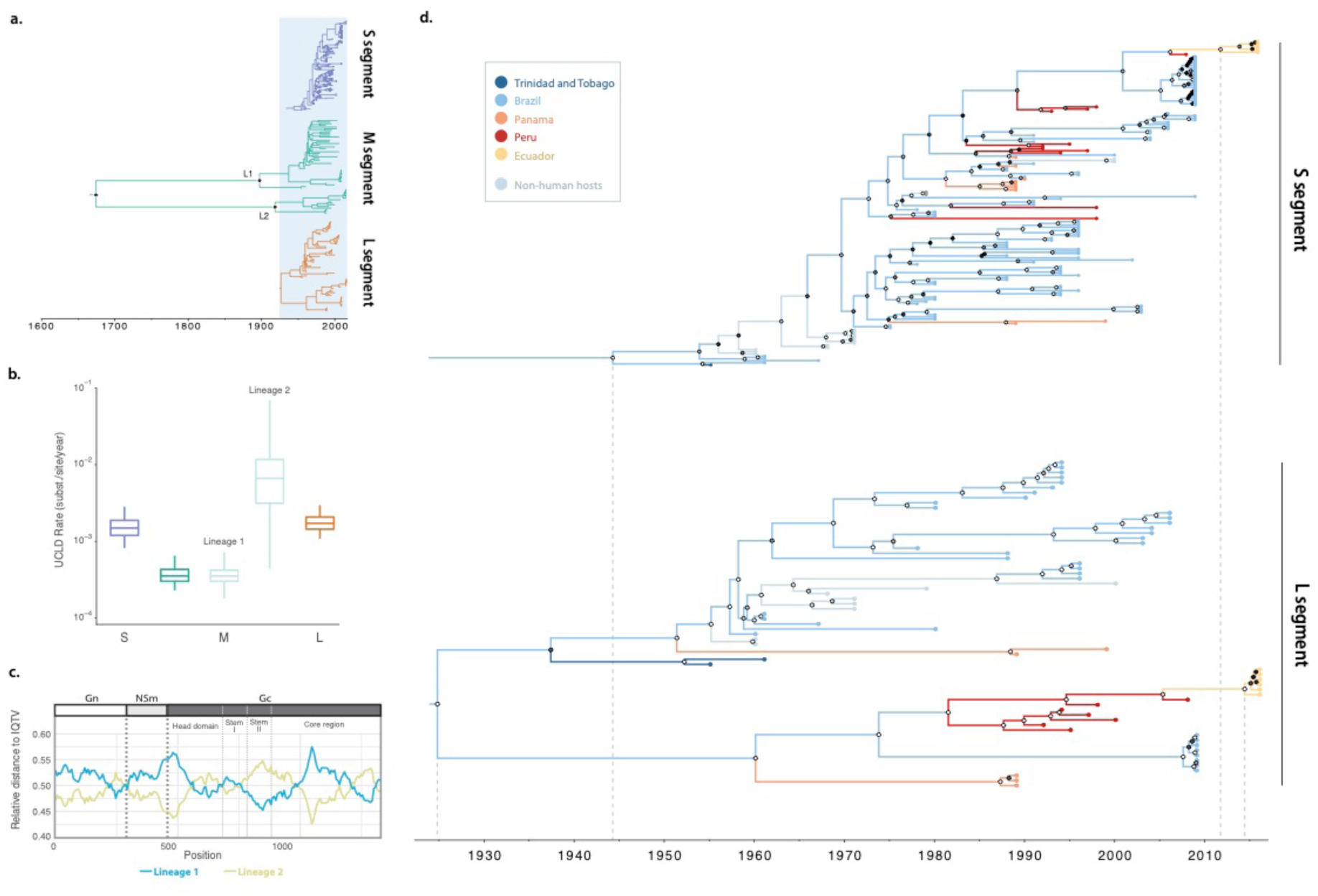
Time-calibrated and evolutionary rate analysis of the OROV genome. **(a)** A comparison of the three segments reveals an older common ancestor for the M segment, resulting in the divergence of two lineages that appear to diversify in the early to mid 1900s (Lineage 1 and Lineage 2, L1 and L2 in the figure). **(b)** The estimated evolutionary rates of the segments reveal a lower rate for the M segment Lineage 1 and a higher rate for Lineage 2. **(c)** A sliding window estimation of the relative genetic distance of each of the two M segment lineages (shown in orange and blue) to Iquitos Virus (IQTV), used as an outgroup. Points of the sequences where the closest genotype to the outgroup changes could be interpreted as recombination events with unsampled sequences. **(d)** An analysis of the younger segments, S and L, reveals emergence date estimates for the virus in the early to mid 20^th^ century, and relatively consistent estimates for the TMRCA of the new Ecuadorian sequences in the early 2010s. The posterior probabilities (PP) of each node is shown based on a greyscale colour scheme (0.0 PP = black to 1.0 PP = white).

In order to explore whether these differences in evolutionary rates among the segment M lineages are a product of rate heterogeneity within the sequences, we estimated the mean relative genetic distance of Lineage 1 and 2 to a closely related outgroup, Iquitos Virus (IQTV), using a sliding window approach. In each window we divided the mean genetic distance of each lineage to the outgroup by the sum of the genetic distances for both lineages, resulting in a *relative* genetic distance to IQTV (see Supplementary R Script). We find that different regions of the M segment display different relative distances to the outgroup, with breakpoints widely distributed within the Gn reading frame, and the core region and head domain of the Gc reading frame. The transition between the Gn and NSm reading frames, and the head and stem domains in the Gc reading frame also correspond approximately to inferred breaking points (Fig. 2c).

In the S segment phylogeny, samples obtained through the 1950s and 1960s are located at the base of the tree. The tree splits into two co-circulating clusters around 1969 (95% HPD: 1964.30-1973.07). Both clusters include sequences mostly from central Brazil and Panama, and one of the clusters suggests multiple introductions to Peru as early as the mid 1970s. The 2009 Mazagão outbreak and the 2016 Ecuador sequences arise from one of these Peruvian clusters, with the latter sharing a common ancestor around late 2011 (95% HPD: 2008.72-2014.88) (Fig. 2d). On the other hand, the L segment tree shows two divergent lineages that split sometime in the early to mid-twentieth century (95% HPDs: 1900.33-1953.31; 1918.02-1978.67) and more closely match the M segment lineage structure. One of these L segment lineages includes most of the Brazilian diversity (the Mazagão sequences are distinct and cluster in the other L segment lineage) and a cluster of sequences from Panama. The other includes a second cluster of Panama sequences and all of the Peruvian diversity (Fig. 2d). The Ecuadorian sequences are again descended from Peruvian strains, with a TMRCA in early 2014 (95% HPD: 2011.34-2015.90). This date is similar in the other segments (the estimated Ecuador TMRCA is mid 2011 in the M segment; 95% HPD: 2007.82-2014.72). These results suggest a recent introduction of OROV to northern Ecuador with no reassortment events leading to its emergence.

### Selection Analyses

We undertook a screen for sites under selection using different models implemented in the HyPhy framework (37). Evidence for both pervasive and episodic selection was found in different genes using different methods (Table 1). Pervasive selection (selection that occurs consistently along the whole phylogeny) was detected in the S segment. Individual sites under pervasive selection were identified in the M segment alignments: for the Gn protein one site was found to be evolving under positive selection (by the SLAC and FEL methods); for the NSm protein one site was identified (by the FEL method); and the Gc protein, two sites were found to be under selection (by the FEL method) (Table 1). Episodic adaptive selection (selection that occurs heterogeneously across the tree branches) was detected in all alignments under the mixed effects model (MEME) (41). In total, MEME identified 6 sites under selection in the RNA dependant RNA polymerase, 7 sites under selection in the M segment polyprotein, and one site under selection in the non-structural N protein. Of these sites, only four were identified by two or more methods: codon 66 of the Gn protein, codon 86 of the NSm protein and codons 269 and 442 of the Gc protein (Table 1). We note that three of these four sites feature more than two alleles, and the frequencies of the most common allele range from 0.44 to 0.73 (Table S1).

**Table 1.** Evidence for adaptive selection on the three genome segments of the OROV genome assessed using different models. Sites identified by more than one method are highlighted in bold.

| Segment | Gene | Length | Pervasive Selection |  |  |  |  |  | Episodic Selection |  |  |  |
| --- | --- | --- | --- | --- | --- | --- | --- | --- | --- | --- | --- | --- |
|  |  |  | *Gene-wise*<br>(*BUSTED*) |  | *Site-wise*<br>(*SLAC*) |  | *Site-wise*<br>(*FEL*) |  | *Site-wise*<br>(*MEME*) |  | *Branch-site*<br>*model (aBSREL)* |  |
|  |  |  | Test<br><0.05 | Mean<br>*dN/dS*<sup>a</sup> | Test<br><0.05 | Sites | Test<br><0.05 | Sites | Test<br><0.05 | Sites | Test<br><0.05 | Sites |
| **S** | N/Ns | 693 nt | YES | 0.071 | NO | -- | NO | -- | YES | 2 | NO | -- |
| **M** | Gn | 918 nt | NO | 0.055 | YES | **66** | YES | **66** | YES | **66** | NO | -- |
|  | NSm | 525 nt | NO | 0.114 | NO | -- | YES | **86** | YES | **86** | NO | -- |
|  | Gc | 2817 nt | NO | 0.050 | NO | -- | YES | **269, 442** | YES | 71, 106,<br>145, **269**,<br>**442**, 645,<br>842, 879 | NO | -- |
| **L** | RdRP | 6756 nt | NO | 0.027 | NO | -- | NO | -- | YES | 237, 1089,<br>1394,<br>1397,<br>2048, 2085 | NO | -- |

<sup>a</sup> Estimated as weighed $\omega$ mean values from the unconstrained model.

### Structural analysis of OROV proteins and sites under selection

Viral glycoproteins are important targets of positive selection and locations of virus adaptation due to their direct interactions with the host immune system. In Orthobunyaviruses, glycoproteins are expressed as a polyprotein, encoded by the three open reading frames of the M segment that represent the Gn, NSm (non-structural) and Gc genes respectively. The main component of the OROV particle spikes is the Gc protein, which is composed of four main domains: a head domain, two stem domains, and a core region (20). We identified eight sites under selection in Gc (four of which fall within the head and stem domains) and two putative, previously unreported N-linked glycosylation sites (Fig. 3a). Estimated dN/dS ratios are similar for each domain (Fig 3b), hence we find no evidence for significantly different selective pressures among domains. However, if dN/dS ratios are estimated separately for the two M segment lineages, then we obtain higher dN/dS estimates for the head and stem domains of Lineage 2, although the large confidence limits mean these differences are not significant (Fig. 3b).

**Figure 3.**
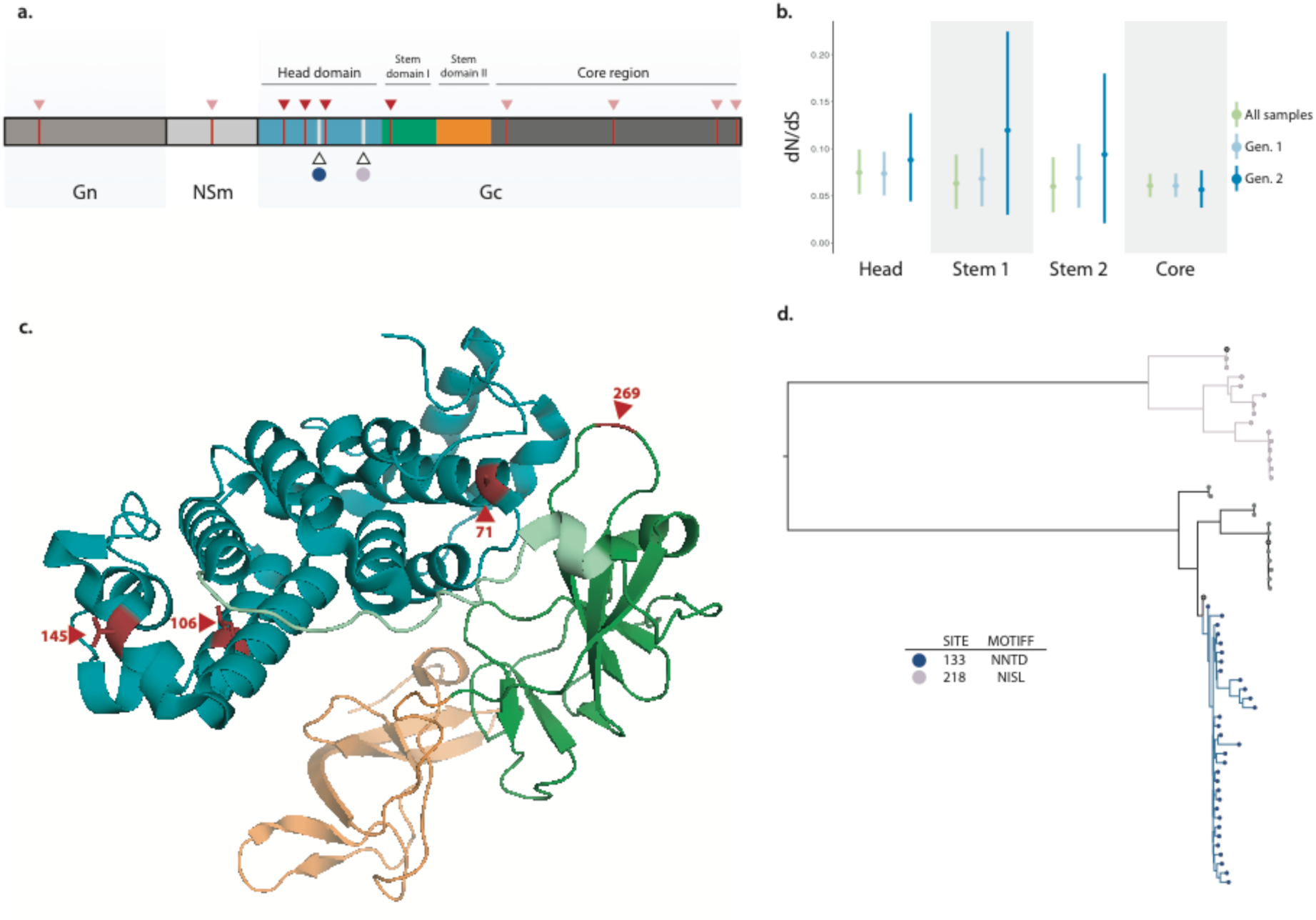
Structure-based mapping of sites identified under selection and previously unreported N-linked glycosylation sites onto the Gc protein. **(a)** A schematic of the OROV M segment polyprotein, which includes the Gn and Gc components of the viral membrane proteins. The Gc protein is subdivided into a head domain (blue), two stalk domains (green and orange), and a C-terminal region, which encodes a putative fusion loop (dark grey). Sites identified as being under positive selection using the MEME approach are shown in red (lighter coloured arrows indicate sites falling outside of the Gc head and stem domains). The two previously undescribed N-glycosylation sites are shown in white. **(b)** Estimated dN/dS ratio of each domain in Gc, obtained for the whole M segment, and for each M segment lineage. **(c)** Sites under positive selection mapped to a composite model generated from previously reported OROV Gc (head domain, PDB ID 6H3X) and SBV Gc (stem domains, PDB ID 6H3S) crystal structures. The OROV head domain is shown in blue, with the C-terminus loop residues shown in light green (taken from the SBV head domain) and SBV stem domains I and II are shown in green and orange, respectively. **(d)** Putative N-linked glycosylation sites identified in this study are mapped onto the phylogeny of the M segment, showing the independent presence the two sites in the two M segment lineages.

We mapped the sites identified as being under positive selection to the available atomic structures of the OROV Gc head domain and the closely related SBV stem domains (Fig 3c). This analysis reveals that three sites under selection (codons 71, 106 and 145) are located at alpha-helixes in the head domain, and one (codon 269) occurs at a loop connecting the stem domain I N-terminus to the head domain (Fig. 3c).

In addition to the sites under positive selection, we identified two previously unreported N-linked glycosylation sequons at residues 133 (motif NNTD) and 218 (motif NISL). Both of these putative glycosylation sites are located on the Gc head domain (Fig. 3a) and do not co-occur together in any of the sequences, supportive of the hypothesise that they may be alternative motifs that could serve similar functional roles, such as shielding antigenic sites. Furthermore, these sites appear to be phylogenetically associated with the different M segment lineages, with motif NNTD being exclusive to Lineage 1 and motif NISL occurring exclusively in Lineage 2 (Fig. 3d). Based on shared ancestry, it is likely that the motif NNTD appeared in a Brazilian common ancestor that excludes the Mazagão outbreak sequences, as this motif is not found in either the Panama or Trinidad sequences belonging to Lineage 1.

Both the putative fusion loop in the C-terminal region of the Gc protein (46)) and two previously reported N-glycosylation sites were conserved amongst all analysed OROV sequences (3).

## Discussion

The evolutionary genomics of OROV in the Americas appears to be driven by a complex combination of factors that are associated with viral life cycle and the nature of the OROV genome. Phylogenetic analysis of the three OROV genome segments reveals different evolutionary histories, with a stronger topological concordance between the medium (M) and long (L) segments, than between those two segments and the S segment. Many of these incongruencies are likely to represent the different evolutionary histories of each segment, arising from reassortment. However, the considerable phylogenetic uncertainty observed for the S phylogeny (in which low bootstrap values and posterior probabilities are observed), limits our ability to draw robust conclusions about tree topology differences involving the S segment. The occurrence of reassortment events appears to be relatively common for OROV in America and may be currently underestimated because (i) there are likely to be many undiscovered OROV lineages and (ii) the virus likely circulates widely in one or more as-yet uncharacterised reservoir populations. Although we found here similarities between the M and L segment phylogenies, previous study of the Bunyamwera group has suggested linkage between the L and S segments (47), and different reassortment patterns have been reported for other Bunyaviruses such as La Crosse virus (LACV) and Snowshoe Hare virus (SSHV) (48). The reassortment of OROV is in line with the observation of naturally occurring reassortant viruses (23, 24), a phenomenon that has been linked to the coinfection of hosts or arthropod vectors. The latter provide a good opportunity for reassortment to occur, as an infected vector can feed on a second vertebrate host infected with a distinct viral strain within a 2-3 day time window and become co-infected (2). On a broader scale, reassortment events could play a crucial role in the evolution of Bunyaviruses in general, and it has been suggested that an increasing number of the recently described species are reassortants (49).

Molecular clock phylogenetic analyses can also be used to infer the emergence times of pathogens and the timescales of outbreaks (50). Some of these tools are particularly applicable to heterochronous data sets, in which sequences from rapidly evolving populations have been sampled longitudinally through time. For molecular clock analyses to be reliable, samples collected at different time points must accumulate sufficient genetic differences to be distinct from one another (51) (i.e. contain sufficient ‘temporal signal’). A combination of factors may affect the accuracy of estimated TMRCAs. Our analysis of the S and L segments suggest that OROV first emerged in the early to mid 1900s, an earlier estimate than previously reported (late 1700s, inferred from the S segment phylogeny) (26). This difference may reflect the fact that our analysis incorporates more sequences, or it may reflect differences in the evolutionary models or prior distributions used in Bayesian phylogenetic inference. It should be also noted that the aforementioned earlier estimate does not report a test for temporal signal in the data set, which could also affect the precision of their estimates.

A key factor of OROV evolutionary dynamics is the distinction between two well supported lineages for the M segment. We estimate that these two lineages split more than two centuries before the estimated TMRCAs of the other two segments. These two lineages are geographically structured in South America, with Lineage 1 representing a central-eastern clade and Lineage 2 representing a western clade. Despite the existence of reassortment, the split between Lineages 1 and 2 is also reflected in the L and S segment phylogenies. Specifically, the L segment shows the same split into two well supported lineages, with the exception of one reassortant (the 2009 Mazagão outbreak sequences; see Fig 2 and S2). The S segment phylogeny is less geographically structured; the ‘western’ Peruvian and Ecuadorian sequences are placed in a single monophyletic clade together with the Mazagão sequences and other Brazilian and Panama sequences, but they show multiple introductions from Brazil to Peru (Fig. 2c). Different evolutionary histories for OROV segments have been previously described for the broader Oropouche virus complex (3, 7); here we show that these differences can be also found but within OROV itself.

The evolutionary rate of the M segment appears to be considerably slower than the remaining segments, an unexpected finding given the tendency of Bunyaviral surface proteins to evolve faster than the nucleocapsid and non-structural proteins (52, 53). Further, when independently estimated, the rate for the western Lineage 2 appears to be an order of magnitude faster than that of Lineage 1. We hypothesise that the low evolutionary rate corresponding to Lineage 1, which matches the overall rate for the M segment, could be an effect of unaccounted recombination events from an unsampled lineage into the observed diversity of the virus (see below).

Next, we sought to explain why the TMRCA of the M segment is substantially older than that of the other two segments, despite the fact that all the segment share overlapping sets of strains, and the fact that the L segment tree shows a similar lineage structure. The age discrepancy between segment TMRCAs suggests that they represent different ancestral viruses, accommodating the notion of widespread reassortment shaping the evolution of OROV. The long internal branches in the M segment phylogeny are unlikely to be artefacts given the temporal signal of the data set (Fig. S1) and therefore suggest that a considerable portion of the genetic diversity remains unsampled. We propose that these results can be understood if the M segment underwent a reassortment event with one such divergent but as yet unsampled lineages that split after the common ancestor of OROV and IQTV (Fig. 4). Hypothetically, the unsampled lineage could represent a distinctly divergent L segment lineage, similar to what is observed for the M segment, which reassorted into an ancestral viral lineage which is represented in the currently available L segment diversity. The reassortment event most likely occurred once, and therefore predates the estimated origin of the recipient M lineage. This would contradict with the younger internal nodes inferred for the L segment; we therefore believe that the true TMRCAs for the M segment lineages are younger than our estimates. The possibility of further recombination of this unsampled lineage into the known sequences suggested in our data set (Fig. 2c) could also affect the precision of our estimates.

**Figure 4.**
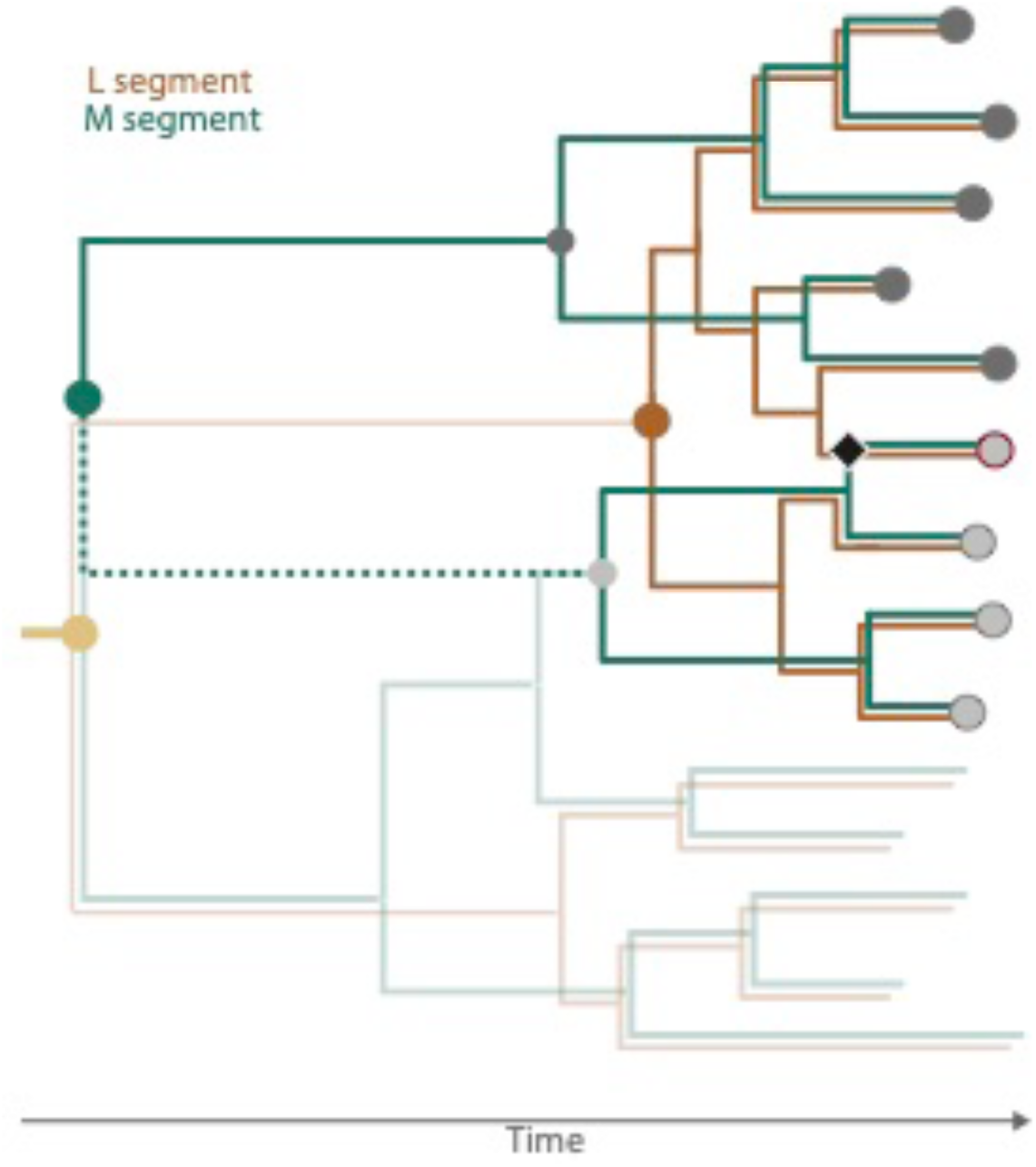
A hypothetic model of the evolution of the OROV genome. The schematic depicts a proposed hypothesis for the evolution of the L (orange branches/outline) and M (green branches/outline) segments. The two M segment lineages (dark and light grey tips) share an older root compared to the L segment, suggesting that the roots of both phylogenies do not represent the same ancestor, and that a more recent reassortment event between the segments has occurred (black diamond) leading to a reassortant clade (red outlined tip). The existence of an unsampled lineage (subdued colours) could represent the unaccounted diversity of the virus that would potentially mirror the two distinct lineages for the M and L segments. Under this model, the long branch inferred for one of the M segment lineages (dotted line) doesn’t represent the true evolutionary history of the virus and fails to account for the proposed reassortment event. The true shared ancestor for both segments (yellow) would probably be as old as the inferred root of the M segment.

The ability of the genome of an emerging virus to adapt to new hosts can affect the likelihood of future outbreaks and is therefore a useful phenomenon to explore. Whilst computational evolutionary analysis can usefully indicate the presence of positive selection and molecular adaptation in virus genomes, sequence data alone cannot resolve the functional and biological context of selected sites. Further, different methods for detecting selection do not always give the same results, hence accuracy and statistical power can be improved by considering the consensus of results generated by multiple models and methods (54). If we combine the results of the multiple tests for positive selection undertaken here, then we identify four sites that are consistently inferred to be under selection (all in the M segment): two sites located on the Gc spike protein, one site on the Gn scaffold protein, and one site on the non-structural NSm protein. Other sites were identified as being under positive selection by only a subset of methods, suggesting that additional regions of the Gc protein undergoing adaptive selection, particularly the head and core domains (Table 1, Fig. 3a).

Three different models indicated the positive selection acting on codon 66 of Gn (Table 1). The N-terminus of closely-related Bunyamwera virus (BUNV) Gn protein is predicted to localise on the outside of the viral particle (55) and assembles as a scaffold to the trimeric Gc spike (19). Codon 66 falls within this predicted extra-membrane domain, potentially exposing it to the host immune system. Furthermore, this protein has been shown to interact closely with the Gc core region to shield the hydrophobic fusion loops and prevent early membrane fusion during viral cell entry. In Phleboviruses such as Rift Valley fever virus (RVFV), the Gn N-terminus folds into a domain that localises on the outermost region of the viral membrane during the pre-fusion, shielded state (56). Codon 66 of the OROV Gn protein could play a role in a similar fusion loop shielding process.

We also detected positive selection on codon 86 of the NSm. The function of this protein among the Orthobunyaviruses is variable, ranging from being essential for viral assembly in BUNV (particularly the N-terminal hydrophobic domains) (57) to being non-essential for viral growth in OROV (7) and other related pathogens such as Maguari virus (MAGV) (58). In RVFV, the NSm protein is non-essential for viral replication (59, 60) but plays a role in the suppression of apoptosis in virus-infected cells (61, 62). Other potential adaptive roles for NSm in different Bunyaviruses have been previously (63), including a role in the infection of arthropod vectors (64). The reason for adaptive evolution in OROV NSm detected here is therefore unclear.

The N-terminal head and stem domains of the orthobunyaviral Gc spike are the only regions for which atomic structures are available (20), making structure-based mapping of sites of selection and N-linked glycosylation feasible. While little is known about the host immune response to OROV, SBV Gc has been shown to be a target of the antibody-mediated immune response following infection (65), and is therefore likely to be under selective pressure by the host antibody response. Codons 269 and 442 (both identified by independent methods as evolving under positive selection) are located on different motifs of the Gc protein, in the first stem domain and core region of the Gc protein, respectively. We mapped codon 269 to a loop on the N-terminus of the stem domain I, which connects to the head domain. Although one cannot discount a potential role in shielding the fusion loop in the core region, crystallographic analysis and comparison of this region with other reported orthobunyavirus Gc structures suggests that this loop may act as a flexible hinge that permits fusogenic rearrangements of the spike complex (20). Furthermore, and in-line with the hypothesis that this region is important in stabilizing the mature spike complex, an identified neutralising antibody against SBV directly interacts with residues closer to the C-terminus of the protein’s head domain, potentially disrupting Gc-mediated trimerization of the spike protein, as presented on viral particles (20). A similar immunogenic mechanism could play a role in the evolution of residue 269, rendering it a potential target of the neutralising antibody response. Of the residues that were identified as evolving under diversifying selection using the MEME approach, codons 106 and 145 are mapped to alpha helixes h8 and h12 and correspond to sites known to interact directly with neutralising antibodies in SBV (20). Thus, these sites could also constitute antibody epitopes and be under immune selection. Furthermore, if this region includes an epitope, the acquisition of N-linked glycosylation at helix h10 may mask the protein surface, similar to the “glycan shields” described for Old World Arenaviruses and HIV-1 (66, 67). The occurrence of non-essential N-linked glycosylation sites affecting viral infectivity and replication has been previously described in BUNV (68) and RVFV (69).

Our results show a complex and dynamic picture of the evolution of OROV in the Americas, shaped by reassortment, spatial structure, and natural selection. Our report incorporates new OROV genome sequences obtained from febrile patients in the province of Esmeraldas in northern Ecuador, representing the first direct detection and whole genome sequencing of the virus in the country (14). Previous prospective studies have suggested that OROV may have been circulating in both the coastal and Amazonian provinces of Ecuador (18, 70), but had failed to directly detect or isolate the virus. Our sequences probably represent an undetected outbreak of OROV on the northern coast of Ecuador, which may also include the southern coast of Colombia (Esmeraldas is close to the Colombian department of Nariño). Our phylogenetic analyses show that the Ecuadorian isolates form a single lineage that is closely related to viruses circulating in Peru, indicating a contiguous expansion of the known area of OROV circulation. The Ecuadorian isolates carry a distinct M segment that is shared with sequences from Peru and Panama, suggesting the presence of a distinct OROV lineage in the western, Pacific-bordering countries of South America.

OROV continues to be a neglected tropical disease and the numbers of genetic sequences for the virus are still limited. The genetic data that are available tend to be biased towards the shorter S segment, which has lower phylogenetic resolution and doesn’t provide a complete picture of OROV evolution. Available sequences are for the most part limited to viral diversity detected during human outbreaks, and therefore fail to capture the dynamics of OROV in reservoir host species and vector populations. The generation of whole OROV genomes, and the broadening of sampling to other countries and host species, are necessary to improve our understanding of the extent and nature of OROV transmission and the importance of reassortment for the evolution of this pathogen.

---

**Figure S1.**
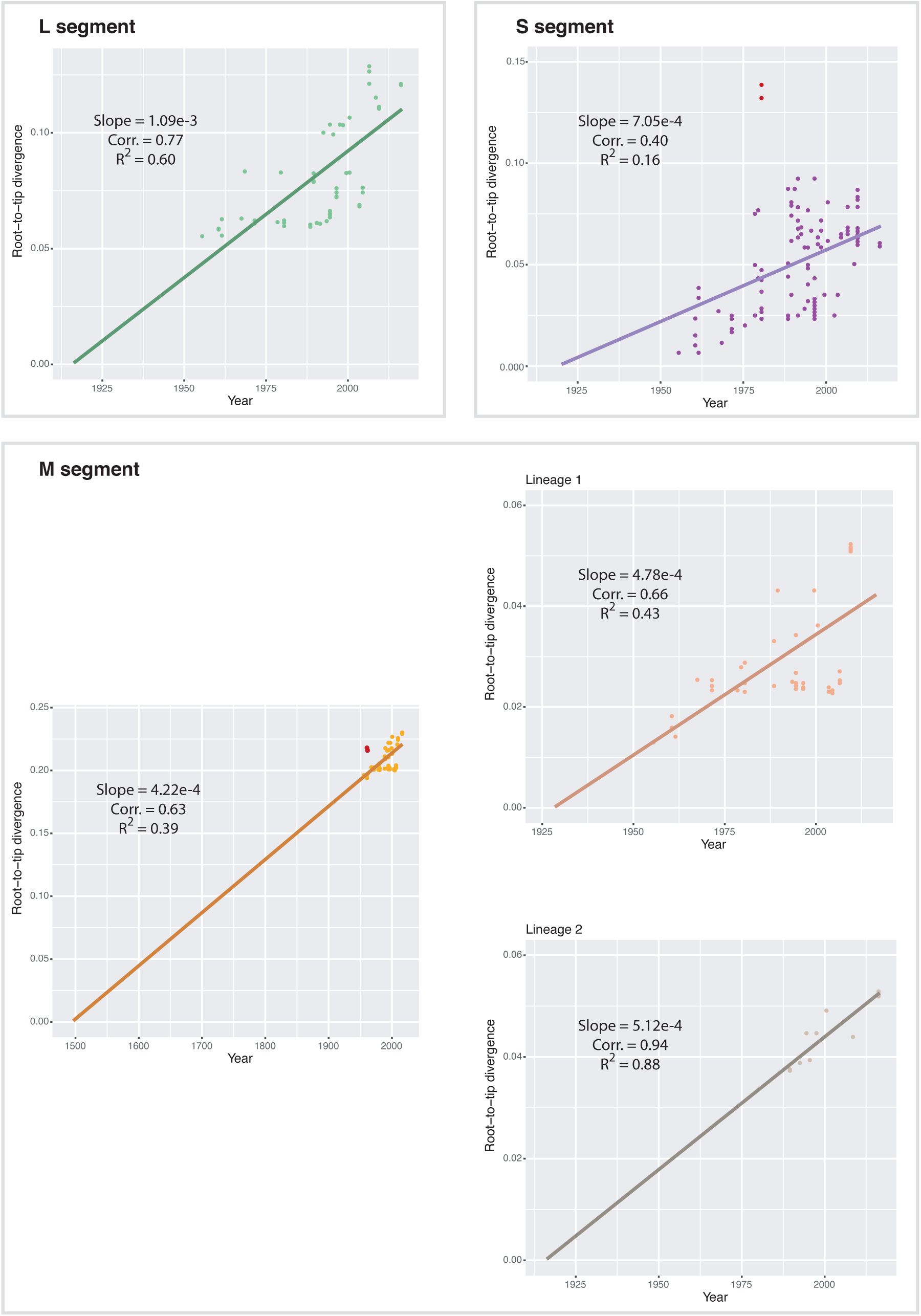
Correlation plots between tip-to-root distances versus sampling estimated in TempEst for the three OROV genome segments. Outliers (determined through visual inspection) are highlighted in red. For the M segment, the complete set of sequences was analysed and visually inspected, and sequences PA01 and PA03 (red) where excluded as outliers prior to the estimation of the individual plots for each lineage. Regression slope (interpreted as rough estimates of the mean evolutionary rate), correlation coefficients and R^2^ values are shown for each plot.

**Figure S2.**
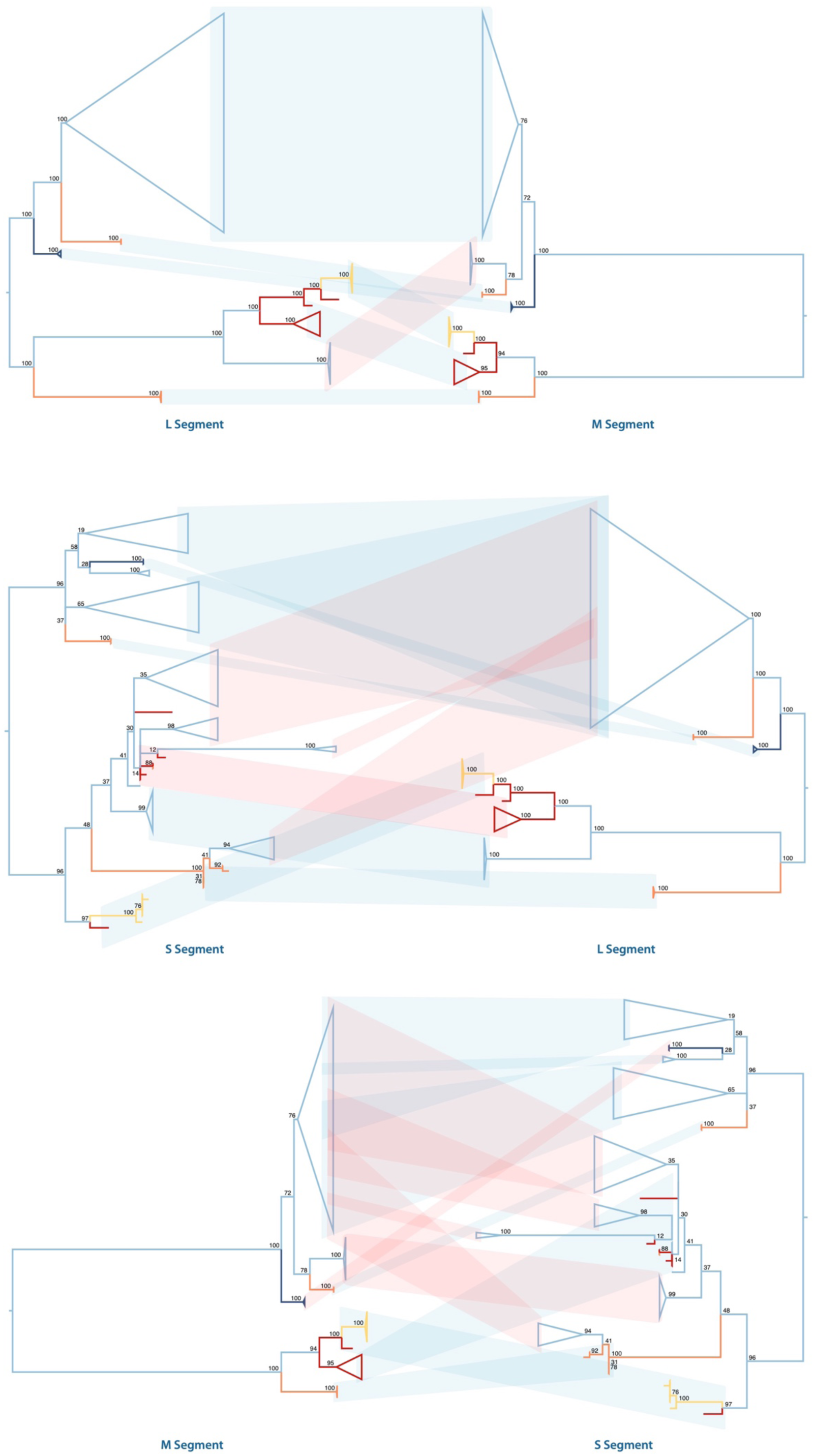
Pairwise comparison of maximum-likelihood phylogenetic trees of the three OROV genome segments suggest widespread reassortment. The L and S trees are midpoint rooted, and the M tree was re-rooted to maximise its topological congruency with its closest match, the L tree. Node support representing 100 bootstraps is shown on the tree nodes, where important monophyletic clades are collapsed in all trees (most collapsed nodes have high node support). Topologically similar clusters are highlighted in blue, while potentially reassortant clusters are highlighted in red.

**Table S1.** Allele frequencies for codons under adaptive selection in the OROV genome. **Supplementary R script** available at GitHub: https://github.com/BernardoGG/OROV_America/blob/master/OROV_SimPlots.R

| Protein | Codon | Identity | Frequency |
| --- | --- | --- | --- |
| Gn | 66 | K | 0.444 |
|  |  | R | 0.333 |
|  |  | A | 0.190 |
|  |  | T | 0.032 |
| NSm | 86 | R | 0.651 |
|  |  | N | 0.206 |
|  |  | S | 0.143 |
| Gc | 269 | D | 0.656 |
|  |  | G | 0.344 |
|  | 442 | T | 0.734 |
|  |  | S | 0.188 |
|  |  | I | 0.078 |

## Notes

https://github.com/BernardoGG/OROV_America

## References

1. Hart TJ, Kohl A, Elliott RM. Role of the NSs protein in the zoonotic capacity of Orthobunyaviruses. Zoonoses Public Health. 2009;56(6-7):285–96.

2. Elliott RM. Orthobunyaviruses: recent genetic and structural insights. Nat Rev Microbiol. 2014;12(10):673–85.

3. Travassos da Rosa JF, de Souza WM, Pinheiro FP, Figueiredo ML, Cardoso JF, Acrani GO, et al. Oropouche Virus: Clinical, Epidemiological, and Molecular Aspects of a Neglected Orthobunyavirus. Am J Trop Med Hyg. 2017;96(5):1019–30.

4. Rodriguez-Morales AJ, Paniz-Mondolfi AE, Villamil-Gomez WE, Navarro JC. Mayaro, Oropouche and Venezuelan Equine Encephalitis viruses: Following in the footsteps of Zika? Travel Med Infect Dis. 2017;15:72–3.

5. Anderson CR, Spence L, Downs WG, Aitken TH. Oropouche virus: a new human disease agent from Trinidad, West Indies. Am J Trop Med Hyg. 1961;10:574–8.

6. Azevedo RS, Nunes MR, Chiang JO, Bensabath G, Vasconcelos HB, Pinto AY, et al. Reemergence of Oropouche fever, northern Brazil. Emerg Infect Dis. 2007;13(6):912–5.

7. Tilston-Lunel NL, Hughes J, Acrani GO, da Silva DE, Azevedo RS, Rodrigues SG, et al. Genetic analysis of members of the species Oropouche virus and identification of a novel M segment sequence. J Gen Virol. 2015;96(Pt 7):1636–50.

8. Vasconcelos HB, Azevedo RS, Casseb SM, Nunes-Neto JP, Chiang JO, Cantuaria PC, et al. Oropouche fever epidemic in Northern Brazil: epidemiology and molecular characterization of isolates. J Clin Virol. 2009;44(2):129–33.

9. Cardoso BF, Serra OP, Heinen LB, Zuchi N, Souza VC, Naveca FG, et al. Detection of Oropouche virus segment S in patients and inCulex quinquefasciatus in the state of Mato Grosso, Brazil. Mem Inst Oswaldo Cruz. 2015;110(6):745–54.

10. da Costa VG, de Rezende Feres VC, Saivish MV, de Lima Gimaque JB, Moreli ML. Silent emergence of Mayaro and Oropouche viruses in humans in Central Brazil. Int J Infect Dis. 2017;62:84–5.

11. Alvarez-Falconi PP, Rios Ruiz BA. [Oropuche fever outbreak in Bagazan, San Martin, Peru: epidemiological evaluation, gastrointestinal and hemorrhagic manifestations]. Rev Gastroenterol Peru. 2010;30(4):334–40.

12. Romero-Alvarez D, Escobar LE. Vegetation loss and the 2016 Oropouche fever outbreak in Peru. Mem Inst Oswaldo Cruz. 2017;112(4):292–8.

13. Watts DM, Phillips I, Callahan JD, Griebenow W, Hyams KC, Hayes CG. Oropouche virus transmission in the Amazon River basin of Peru. Am J Trop Med Hyg. 1997;56(2):148–52.

14. Wise EL, Pullan ST, Marquez S, Paz V, Mosquera JD, Zapata S, et al. Isolation of Oropouche Virus from Febrile Patient, Ecuador. Emerg Infect Dis. 2018;24(5):935–7.

15. Nunes MR, Martins LC, Rodrigues SG, Chiang JO, Azevedo Rdo S, da Rosa AP, et al. Oropouche virus isolation, southeast Brazil. Emerg Infect Dis. 2005;11(10):1610–3.

16. Romero-Alvarez D, Escobar LE. Oropouche fever, an emergent disease from the Americas. Microbes Infect. 2018;20(3):135–46.

17. Sakkas H, Bozidis P, Franks A, Papadopoulou C. Oropouche Fever: A Review. Viruses. 2018;10(4).

18. Manock SR, Jacobsen KH, de Bravo NB, Russell KL, Negrete M, Olson JG, et al. Etiology of acute undifferentiated febrile illness in the Amazon basin of Ecuador. Am J Trop Med Hyg. 2009;81(1):146–51.

19. Bowden TA, Bitto D, McLees A, Yeromonahos C, Elliott RM, Huiskonen JT. Orthobunyavirus ultrastructure and the curious tripodal glycoprotein spike. PLoS Pathog. 2013;9(5):e1003374.

20. Hellert J, Aebischer A, Wernike K, Haouz A, Brocchi E, Reiche S, et al. Orthobunyavirus spike architecture and recognition by neutralizing antibodies. Nat Commun. 2019;10(1):879.

21. Briese T, Bird B, Kapoor V, Nichol ST, Lipkin WI. Batai and Ngari viruses: M segment reassortment and association with severe febrile disease outbreaks in East Africa. J Virol. 2006;80(11):5627–30.

22. Saeed MF, Wang H, Suderman M, Beasley DW, Travassos da Rosa A, Li L, et al. Jatobal virus is a reassortant containing the small RNA of Oropouche virus. Virus Res. 2001;77(1):25–30.

23. Aguilar PV, Barrett AD, Saeed MF, Watts DM, Russell K, Guevara C, et al. Iquitos virus: a novel reassortant Orthobunyavirus associated with human illness in Peru. PLoS Negl Trop Dis. 2011;5(9):e1315.

24. Navarro JC, Giambalvo D, Hernandez R, Auguste AJ, Tesh RB, Weaver SC, et al. Isolation of Madre de Dios Virus (Orthobunyavirus; Bunyaviridae), an Oropouche Virus Species Reassortant, from a Monkey in Venezuela. Am J Trop Med Hyg. 2016;95(2):328–38.

25. Bernardes-Terzian AC, de-Moraes-Bronzoni RV, Drumond BP, Da Silva-Nunes M, da-Silva NS, Urbano-Ferreira M, et al. Sporadic oropouche virus infection, acre, Brazil. Emerg Infect Dis. 2009;15(2):348–50.

26. Vasconcelos HB, Nunes MR, Casseb LM, Carvalho VL, Pinto da Silva EV, Silva M, et al. Molecular epidemiology of Oropouche virus, Brazil. Emerg Infect Dis. 2011;17(5):800–6.

27. Greninger AL, Naccache SN, Federman S, Yu G, Mbala P, Bres V, et al. Rapid metagenomic identification of viral pathogens in clinical samples by real-time nanopore sequencing analysis. Genome Med. 2015;7:99.

28. Li H, Durbin R. Fast and accurate long-read alignment with Burrows-Wheeler transform. Bioinformatics. 2010;26(5):589–95.

29. Penedos AR, Myers R, Hadef B, Aladin F, Brown KE. Assessment of the Utility of Whole Genome Sequencing of Measles Virus in the Characterisation of Outbreaks. PLoS One. 2015;10(11):e0143081.

30. Kearse M, Moir R, Wilson A, Stones-Havas S, Cheung M, Sturrock S, et al. Geneious Basic: an integrated and extendable desktop software platform for the organization and analysis of sequence data. Bioinformatics. 2012;28(12):1647–9.

31. Martin DP, Murrell B, Golden M, Khoosal A, Muhire B. RDP4: Detection and analysis of recombination patterns in virus genomes. Virus Evol. 2015;1(1):vev003.

32. Stamatakis A. RAxML version 8: a tool for phylogenetic analysis and post-analysis of large phylogenies. Bioinformatics. 2014;30(9):1312–3.

33. Rambaut A, Lam TT, Max Carvalho L, Pybus OG. Exploring the temporal structure of heterochronous sequences using TempEst (formerly Path-O-Gen). Virus Evol. 2016;2(1):vew007.

34. Suchard MA, Lemey P, Baele G, Ayres DL, Drummond AJ, Rambaut A. Bayesian phylogenetic and phylodynamic data integration using BEAST 1.10. Virus Evol. 2018;4(1):vey016.

35. Drummond AJ, Rambaut A, Shapiro B, Pybus OG. Bayesian coalescent inference of past population dynamics from molecular sequences. Mol Biol Evol. 2005;22(5):1185–92.

36. Drummond AJ, Ho SY, Phillips MJ, Rambaut A. Relaxed phylogenetics and dating with confidence. PLoS Biol. 2006;4(5):e88.

37. Pond SL, Frost SD, Muse SV. HyPhy: hypothesis testing using phylogenies. Bioinformatics. 2005;21(5):676–9.

38. Weaver S, Shank SD, Spielman SJ, Li M, Muse SV, Kosakovsky Pond SL. Datamonkey 2.0: a modern web application for characterizing selective and other evolutionary processes. Mol Biol Evol. 2018.

39. Murrell B, Weaver S, Smith MD, Wertheim JO, Murrell S, Aylward A, et al. Gene-wide identification of episodic selection. Mol Biol Evol. 2015;32(5):1365–71.

40. Kosakovsky Pond SL, Frost SD. Not so different after all: a comparison of methods for detecting amino acid sites under selection. Mol Biol Evol. 2005;22(5):1208–22.

41. Murrell B, Wertheim JO, Moola S, Weighill T, Scheffler K, Kosakovsky Pond SL. Detecting individual sites subject to episodic diversifying selection. PLoS Genet. 2012;8(7):e1002764.

42. Smith MD, Wertheim JO, Weaver S, Murrell B, Scheffler K, Kosakovsky Pond SL. Less is more: an adaptive branch-site random effects model for efficient detection of episodic diversifying selection. Mol Biol Evol. 2015;32(5):1342–53.

43. Schrodinger, LLC. The PyMOL Molecular Graphics System, Version 1.8. 2015.

44. Lemey P, Minin VN, Bielejec F, Kosakovsky Pond SL, Suchard MA. A counting renaissance: combining stochastic mapping and empirical Bayes to quickly detect amino acid sites under positive selection. Bioinformatics. 2012;28(24):3248–56.

45. Nunes MRT, de Souza WM, Savji N, Figueiredo ML, Cardoso JF, da Silva SP, et al. Oropouche orthobunyavirus: Genetic characterization of full-length genomes and development of molecular methods to discriminate natural reassortments. Infect Genet Evol. 2019;68:16–22.

46. Harrison SC. Viral membrane fusion. Virology. 2015;479–480:498-507.

47. Iroegbu CU, Pringle CR. Genetic interactions among viruses of the Bunyamwera complex. J Virol. 1981;37(1):383–94.

48. Urquidi V, Bishop DH. Non-random reassortment between the tripartite RNA genomes of La Crosse and snowshoe hare viruses. J Gen Virol. 1992;73 (Pt 9):2255–65.

49. Briese T, Calisher CH, Higgs S. Viruses of the family Bunyaviridae: are all available isolates reassortants? Virology. 2013;446(1-2):207–16.

50. Wohl S, Schaffner SF, Sabeti PC. Genomic Analysis of Viral Outbreaks. Annu Rev Virol. 2016;3(1):173–95.

51. Drummond AJ, Pybus OG, Rambaut A, Forsberg R, Rodrigo AG. Measurably evolving populations. Trends in Ecology & Evolution. 2003;18(9):481–8.

52. Freire CC, Iamarino A, Soumare PO, Faye O, Sall AA, Zanotto PM. Reassortment and distinct evolutionary dynamics of Rift Valley Fever virus genomic segments. Sci Rep. 2015;5:11353.

53. Lam TT, Liu W, Bowden TA, Cui N, Zhuang L, Liu K, et al. Evolutionary and molecular analysis of the emergent severe fever with thrombocytopenia syndrome virus. Epidemics. 2013;5(1):1–10.

54. Anisimova M, Bielawski JP, Yang Z. Accuracy and power of bayes prediction of amino acid sites under positive selection. Mol Biol Evol. 2002;19(6):950–8.

55. Shi X, Botting CH, Li P, Niglas M, Brennan B, Shirran SL, et al. Bunyamwera orthobunyavirus glycoprotein precursor is processed by cellular signal peptidase and signal peptide peptidase. Proc Natl Acad Sci U S A. 2016;113(31):8825–30.

56. Halldorsson S, Li S, Li M, Harlos K, Bowden TA, Huiskonen JT. Shielding and activation of a viral membrane fusion protein. Nat Commun. 2018;9(1):349.

57. Shi X, Kohl A, Leonard VH, Li P, McLees A, Elliott RM. Requirement of the N-terminal region of orthobunyavirus nonstructural protein NSm for virus assembly and morphogenesis. J Virol. 2006;80(16):8089–99.

58. Pollitt E, Zhao J, Muscat P, Elliott RM. Characterization of Maguari orthobunyavirus mutants suggests the nonstructural protein NSm is not essential for growth in tissue culture. Virology. 2006;348(1):224–32.

59. Gerrard SR, Bird BH, Albarino CG, Nichol ST. The NSm proteins of Rift Valley fever virus are dispensable for maturation, replication and infection. Virology. 2007;359(2):459–65.

60. Won S, Ikegami T, Peters CJ, Makino S. NSm and 78-kilodalton proteins of Rift Valley fever virus are nonessential for viral replication in cell culture. J Virol. 2006;80(16):8274–8.

61. Won S, Ikegami T, Peters CJ, Makino S. NSm protein of Rift Valley fever virus suppresses virus-induced apoptosis. J Virol. 2007;81(24):13335–45.

62. Terasaki K, Won S, Makino S. The C-terminal region of Rift Valley fever virus NSm protein targets the protein to the mitochondrial outer membrane and exerts antiapoptotic function. J Virol. 2013;87(1):676–82.

63. Eifan S, Schnettler E, Dietrich I, Kohl A, Blomstrom AL. Non-structural proteins of arthropod-borne bunyaviruses: roles and functions. Viruses. 2013;5(10):2447–68.

64. Crabtree MB, Kent Crockett RJ, Bird BH, Nichol ST, Erickson BR, Biggerstaff BJ, et al. Infection and transmission of Rift Valley fever viruses lacking the NSs and/or NSm genes in mosquitoes: potential role for NSm in mosquito infection. PLoS Negl Trop Dis. 2012;6(5):e1639.

65. Wernike K, Aebischer A, Roman-Sosa G, Beer M. The N-terminal domain of Schmallenberg virus envelope protein Gc is highly immunogenic and can provide protection from infection. Sci Rep. 2017;7:42500.

66. Watanabe Y, Raghwani J, Allen JD, Seabright GE, Li S, Moser F, et al. Structure of the Lassa virus glycan shield provides a model for immunological resistance. Proc Natl Acad Sci U S A. 2018;115(28):7320–5.

67. Seabright GE, Doores KJ, Burton DR, Crispin M. Protein and Glycan Mimicry in HIV Vaccine Design. J Mol Biol. 2019;431(12):2223–47.

68. Shi X, Brauburger K, Elliott RM. Role of N-linked glycans on bunyamwera virus glycoproteins in intracellular trafficking, protein folding, and virus infectivity. J Virol. 2005;79(21):13725–34.

69. Phoenix I, Nishiyama S, Lokugamage N, Hill TE, Huante MB, Slack OA, et al. N-Glycans on the Rift Valley Fever Virus Envelope Glycoproteins Gn and Gc Redundantly Support Viral Infection via DC-SIGN. Viruses. 2016;8(5).

70. Forshey BM, Guevara C, Laguna-Torres VA, Cespedes M, Vargas J, Gianella A, et al. Arboviral etiologies of acute febrile illnesses in Western South America, 2000-2007. PLoS Negl Trop Dis. 2010;4(8):e787.

