## Supplementary figures and images for "The evolutionary dynamics of Oropouche Virus (OROV) in South America"

### Supplementary Figure 1

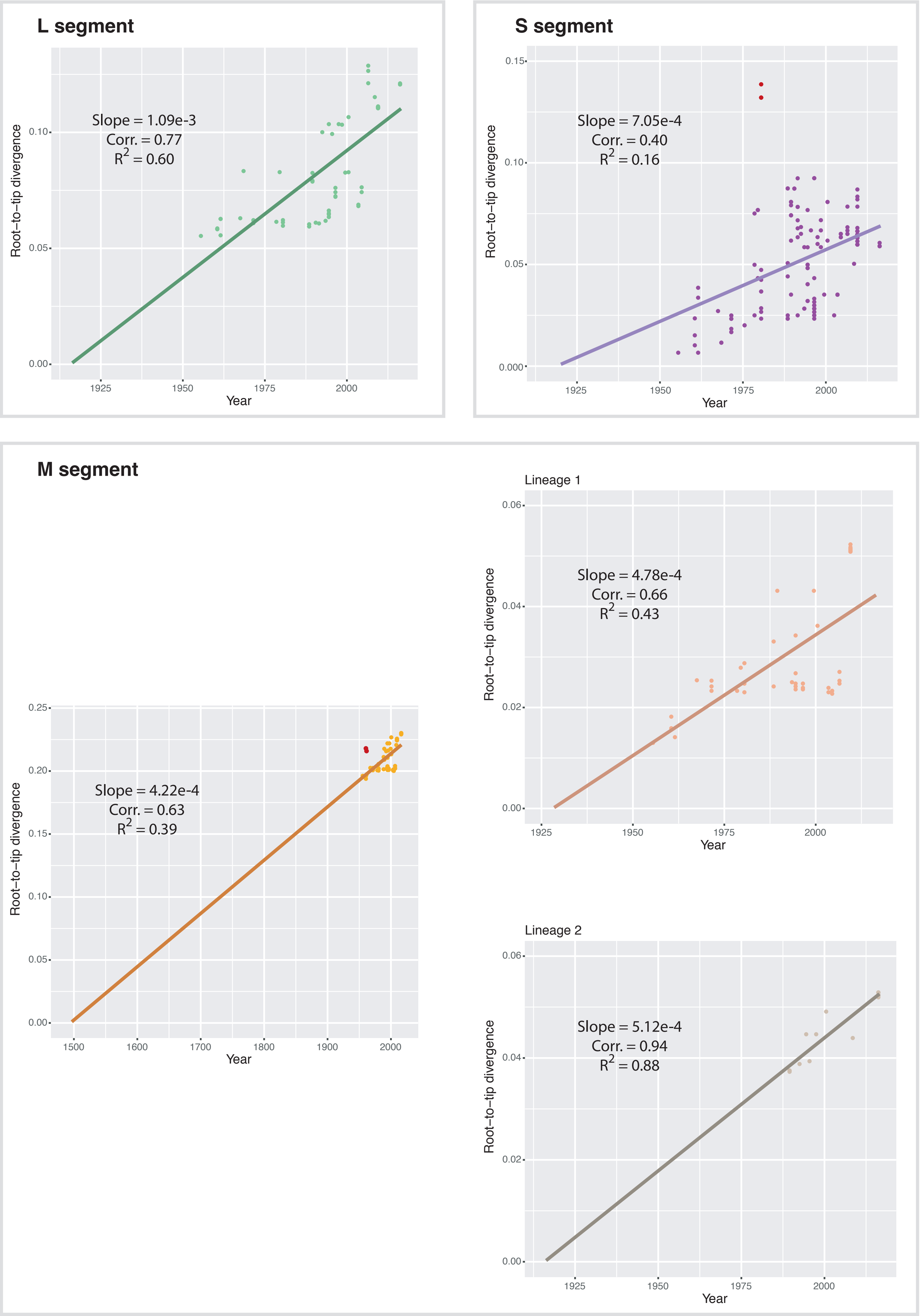

### Supplementary Figure 2

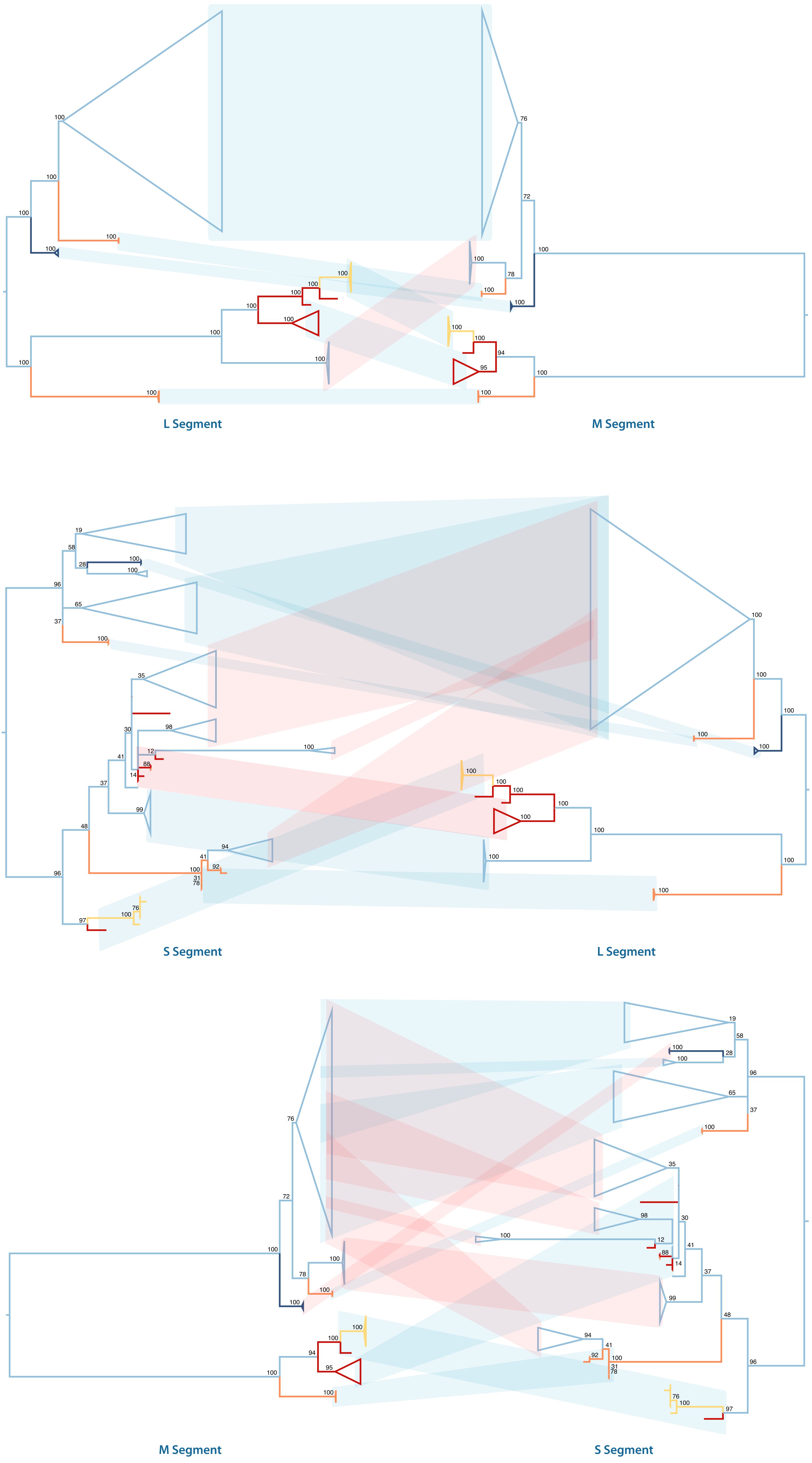
